# *SLC6A4* binding site and acute prosocial effects of (+/-)-3,4-methylendioxymethamphetamine (MDMA) are evolutionarily conserved in *Octopus bimaculoides*

**DOI:** 10.1101/301192

**Authors:** Eric Edsinger, Gül Dölen

**Affiliations:** Josephine Bay Paul Center for Comparative Molecular Biology and Evolution, Marine Biological Laboratory; Department of Neuroscience, Brain Science Institute, Wendy Klag Institute, Kavli Neuroscience Discovery Institute, Johns Hopkins University, School of Medicine

**Author notes:** Correspondence to: Gül Dölen, Department of Neuroscience and Brain Science Institute, 855 N. Wolfe St., Rangos Bldg. Rm. 286, Johns Hopkins University School of Medicine, Baltimore, MD.; E.

## Abstract

Human and octopus lineages are separated by over 500 million years of evolution, and show divergent anatomical patterns of brain organization. Moreover, while humans exhibit highly complex social behaviors, octopuses are thought to be largely asocial and solitary. Despite these differences, growing evidence suggests that ancient neurotransmitter systems are shared across vertebrate and invertebrate species, and in many cases enable overlapping functions. Here we provide evidence that, as in humans, the atypical amphetamine derivative (+/-)-3,4-methylendioxymethamphetamine (MDMA) enhances acute prosocial behaviors in *Octopus bimaculoides.* This finding is paralleled by the evolutionary conservation of the serotonin transporter (SERT, encoded by the *Slc6A4* gene) binding site of MDMA in the *O. bimaculoides* genome. Taken together, these data provide evidence that the neural mechanisms subserving social behaviors exist in *O. bimaculoides*, and indicate that the role of serotonergic neurotransmission in regulating social behaviors is evolutionarily conserved.

**ONE SENTENCE SUMMARY:** Here we provide evidence that the atypical amphetamine derivative (+/-)-3,4-methylendioxymethamphetamine (MDMA) increases acute social approach behaviors in *Octopus bimaculoides*, a finding that is paralleled by the evolutionary conservation of the SLC6A4 binding site of MDMA.

## MAIN TEXT

Sociality is widespread across the animal kingdom, with numerous examples in both invertebrate (e.g. bees, ants, termites, and shrimps) and vertebrate (e.g. fishes, birds, rodents, and primates) lineages (*1*). Serotonin is an evolutionarily ancient molecule (*2*), which has been implicated in regulating both invertebrate (*3*) and vertebrate (*4*) social behaviors, raising the possibility that this neurotransmitter’s prosocial functions may be conserved across evolution. Members of the order *Octopoda* are predominantly asocial and solitary (*5*). Although at this time it is unknown whether serotonergic signaling systems are functionally conserved in octopuses, ethological studies indicate that agonistic behaviors are suspended during mating (*6*–*8*), suggesting that neural mechanisms subserving social behaviors exist in octopuses, but are suppressed outside the reproductive period.

In order to experimentally quantify octopus social behaviors, here we adapted the 3-chambered social approach assay routinely used in rodents (*9*, *10*) for use in *Octopus bimaculoides* (*11*, *12*). As diagramed in **Fig. 1A**, a glass aquarium partitioned into three equally sized chambers containing a novel object or a novel conspecific (male or female social object) in each of the lateral chambers, and an empty center chamber, served as the test arena. The amount of time subject animals spent freely exploring each chamber was recorded during 30-minute test sessions. As shown in **Fig. 1B**, when the social object was a female, male and female subject animals spent significantly more time in the social chamber compared to the center chamber, whereas when the social object was a male, subject animals spent significantly more time in the object chamber compared to the center chamber. Comparisons between conditions revealed that the time spent with the novel object was significantly increased (**Fig. 1C and D**) while time spent with the social object was significantly decreased (**Fig. 1G and H**) when the social object was a male versus a female. The time spent in the center chamber was unchanged across conditions (**Fig. 1E and F**). In order to determine the contribution of social novelty to social approach behaviors, subject animals were next tested on a 2-phase variant of the 3-chambered social approach task (*10*). As shown in **Fig. 1I**, during phase I, lateral chambers contained either a novel object or a novel social object. In phase II, the novel object was replaced with a novel social object (**Fig. 1J**). The sex of the social object was counterbalanced such that equal numbers of males and females were presented as the social object in phase I, and in phase II opposite sexes were used as familiar and novel social objects. As shown in **Fig. 1K**, under these conditions, the amount of time spent was not significantly different across chambers, indicating that social novelty is not a superseding factor in determining approach towards a male or female social object. These data provide the first quantification of social approach behavior in *O. bimaculoides*, and demonstrate a significant preference for interactions with female versus male social objects, which is not modified by social novelty.

**Fig. 1.**
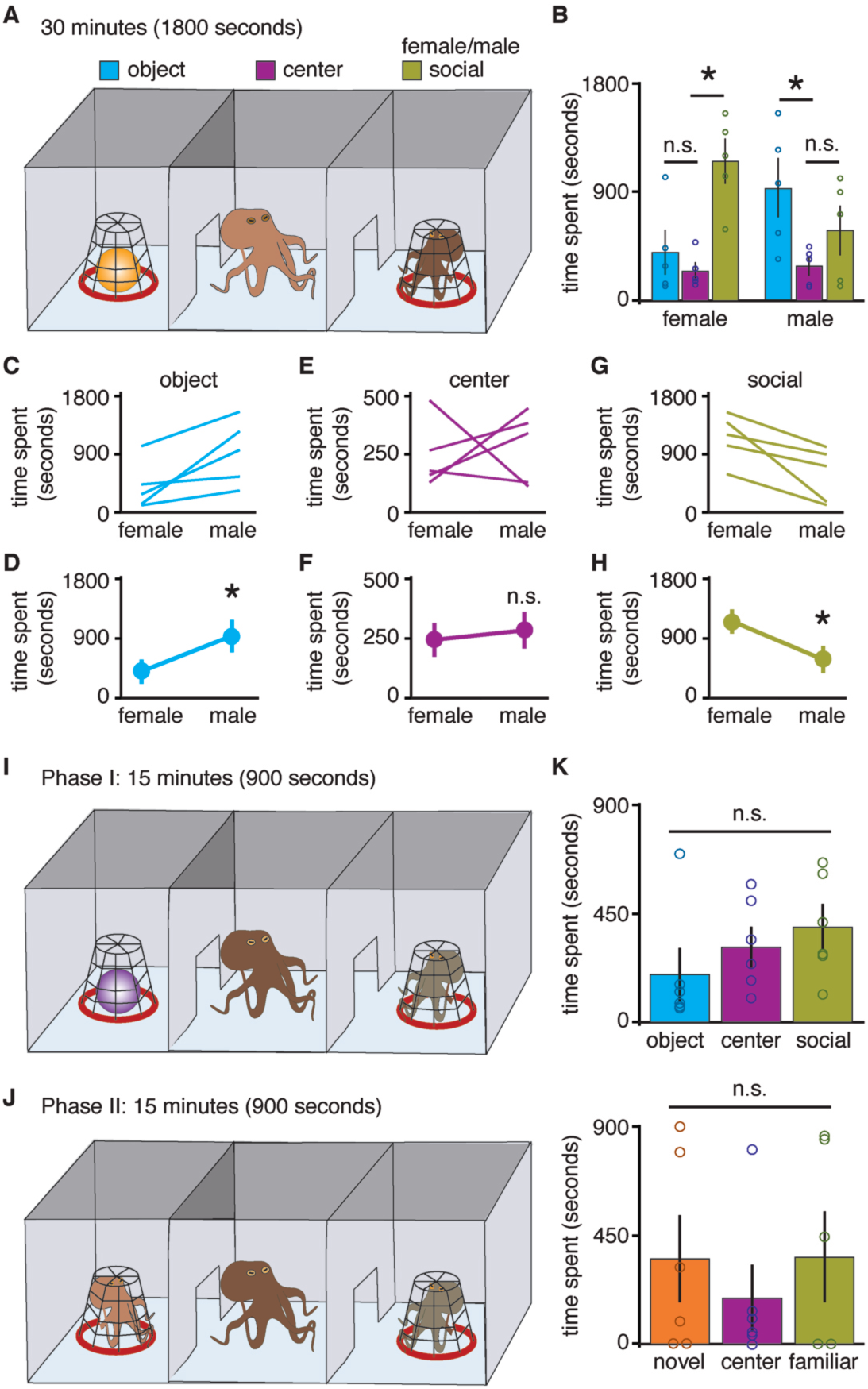
Social approach behaviors towards male and female conspecifics in *Octopus bimaculoides*. **A.** Diagram illustrating three-chambered social approach assay. **B**. Quantification of time spent in each chambers during 30-minute test sessions (n = 5, Two-way repeated measures ANOVA: P = 0.0110; post-hoc unpaired t-test female social object, social versus center P = 0.0009, object versus center P = 0.4031; male social object, social versus center P = 0.1710, object versus center P = 0.0224). **C-H**. Comparisons between female versus male social object conditions for each chamber. Paired t-test female versus male: novel object chamber P = 0.0386; social object chamber P = 0.0281; center chamber P = 0.7544). **I-J.** Diagrams illustrating the two-phase variant of the three-chambered social approach task. During phase I (I), lateral chambers contained either a novel object or a novel social object. In phase II, the novel object was replaced with a novel social object (**J**). **K**. Quantification of time spent in each chambers during 15-minute test sessions, showed no significant effect of novelty on social approach behaviors (n = 6, Two-way repeated measures ANOVA: P = 0.5684).

Recently we completed whole genome sequencing and assembly in *O. bimaculoides (IS)*. Because octopuses are thought to be the most behaviorally advanced invertebrates, this resource enables testing of molecular homology for complex behaviors, despite anatomical differences in brain organization across evolutionarily distant lineages (*14*). In this context, it is interesting to note that in vertebrates (humans and rodents), the atypical amphetamine derivative (+/-)-3,4-methylendioxymethamphetamine (MDMA) is known for its powerful prosocial properties (*15*–*17*). Furthermore, the sixth trans-membrane domain (TM6) of the serotonin transporter (SERT), encoded by the *SLC6A4* gene (*18*), has been identified as the principle binding site of MDMA (*19*–*21*). To determine if this binding site is conserved in *O. bimaculoides*, we performed a molecular phylogenetic analysis of the solute carrier (SLC) 6A subfamily of neurotransmitter transporter proteins (*22*) across selected diverse taxa (**Tables S1 and S2**). Blasting a reference gene set against twenty-one proteomes resulted in a final set of 503 sequences. These sequences were used to generate a maximum likelihood phylogenetic tree (RAxML’s “best tree”) for SLC6A across all twenty-one species (**Fig. 2A**), along with 156 bootstrap trees and a bootstrap “consensus tree” (**Fig. S1**). The “best tree” included the full set of human SLC6A genes and no additional human genes from other SLC families. Expected SLC6A4 family members were found for fruit fly (*Drosophila melanogaster*), worm (*Caenorhabditis elegans*), and all vertebrate species tested, although the “best tree” (**Fig. 2B**) and the “consensus tree” (**Fig. S1**) differed slightly for worm. Surprisingly, two copies of SLC6A4 were found in molluscs, including *O. bimaculoides.* One copy in octopus, named here *Slc6a4*-(*1*) (protein Ocbimv22009795m.p; gene model Ocbimv22008529m.g) was part of a clade that contained only molluscs. The second copy, named here *Slc6a4*-(*2*) (protein Ocbimv22009795m.p; gene model Ocbimv22009795m.g), was in a group that included diverse species, like fly and worm, but no mammals, vertebrates, or other deuterostomes. It is unclear if this duplication is a molluscan innovation, or has more ancient origins in the Lophotrochozoa. Two and three copies of SLC6A4 were present in zebrafish (*Danio rerio*) and zebra finch (*Taeniopygia guttata*), respectively, but only a single copy was present in mammals. These patterns may reflect differential loss of SLC6A4 following two rounds of vertebrate genome duplication (*23*).

**Fig. 2.**
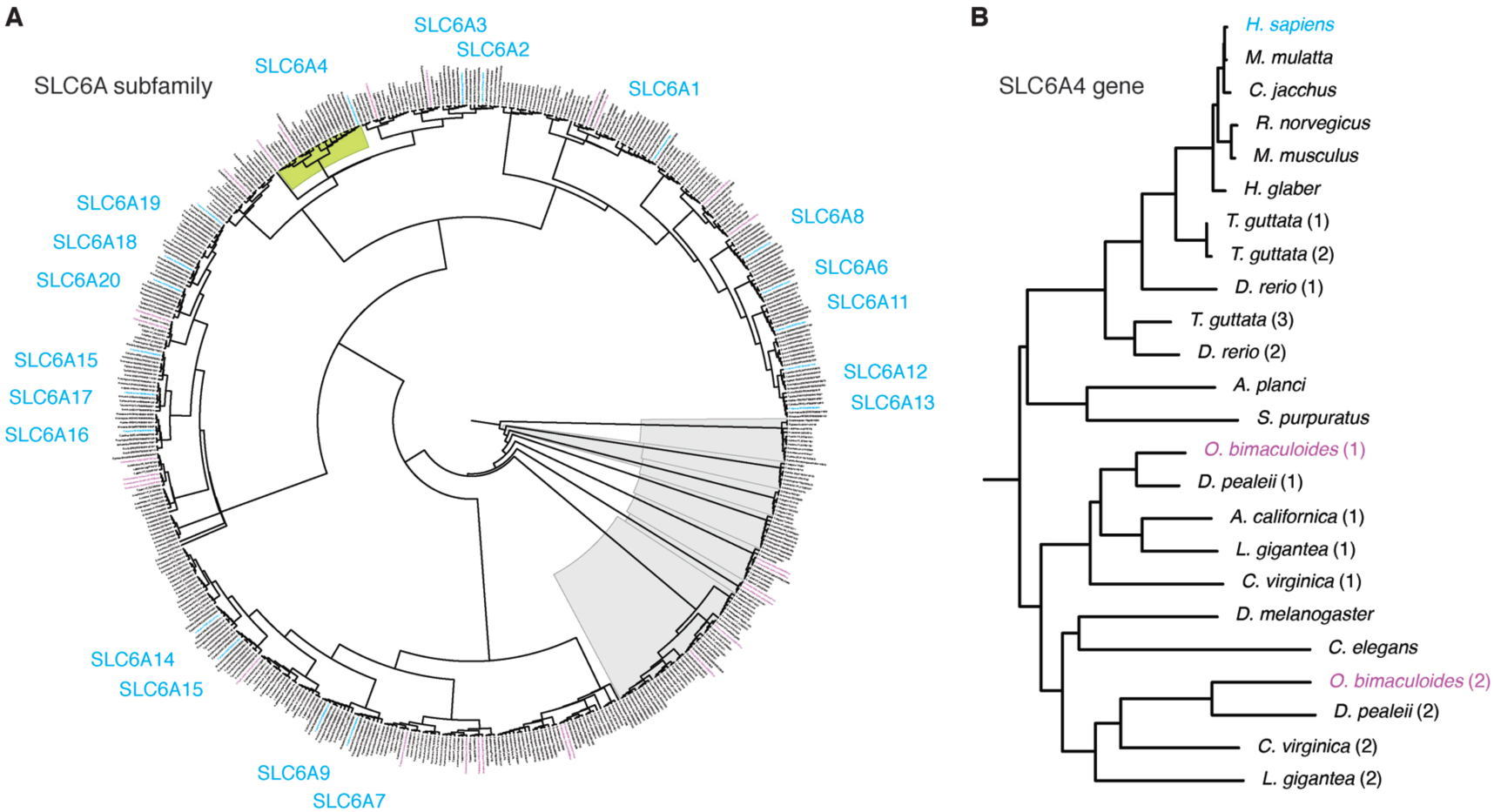
Phylogenetic trees of SLC6A and SLC6A4 gene families. **A-B.** Maximum likelihood trees of (A) SLC6A transporters and (B) SLC6A4 serotonin transporters in select taxa. Species are mapped to tree and protein identifiers in **Table S3**. For a larger version of panel A, see ****Fig. S6****. **A**. A maximum likelihood “best tree” for the SLC6A gene family. The maximum likelihood tree produced by RAxML includes 503 proteins and 21 species, with tree building based on a MAFFT alignment of full-length sequences. **B**. The SLC6A4 gene family, a subtree of the maximum likelihood “best tree” in ****Fig. 1A****.

In contrast to octopus, honey bee (*Apis mellifera*), leaf cutter ant (*Atta cephalotes*), spider (*Stegodyphus mimosarum*), and anemone (*Nematostella vectensis*) all lacked SLC6A4 orthologs. To understand these losses, we next examined all monoamine transporters, including human SLC6A2 (DAT), SLC6A3 (NET), and SLC6A4 (SERT), along with their outgroup, a branch that includes the fruit fly inebriated (*Ine*) gene (**Fig. S2**). Anemone had multiple paralogs within the *Ine* clade. Zebrafish and more basal-branching deuterostomes were also present, but vertebrates, including human, were absent. In contrast, anemone was absent in SLC64A and throughout the monoamine transporter clade. Thus, monoamine transporters may represent an ancient innovation that arose early in bilaterian evolution. Honey bee, leaf cutter ant, fruit fly, and worm all had one or more monoamine transporter proteins outside the SLC6A4 family, while only fruit fly and worm had proteins within the family, supporting the conclusion that SLC6A4 has been lost in hymenopteran insects (ants, bees, wasps, and sawflies). Taken together, these studies underscore the complexity of monoamine transporter evolution in animals, and identify clear orthologs of human SLC6A4 in octopus.

The binding pocket of SLC6A4 is formed by a subset of twelve transmembrane domains, including TM6 (*18*). Previous studies have revealed an especially important role for the region of TM6 that spans amino acids 333 to 336 (indicated in green in **Fig. 3A**), as it provides an overlapping binding pocket for MDMA and serotonin (*19*). Furthermore, residue Ser336 has been implicated in the MDMA induced conformational change not observed with serotonin (*19*). Significantly, for both octopus orthologues within this region, there is 100% percent identity when aligned to human SLC6A4 (**Figs. 3; Figs. S3, and S4**). More generally, many of the domains forming the binding pocket are highly conserved in comparison to other domains and to the full-length protein (**Fig. S4**). For instance, TM6 in Slc6a4-(1) has 95.7% percent identity to human, compared to 53.4% for the protein, and TM8 in Slc6a4-(2) has 91.0% percent identity, compared to 39.0% for the protein (**Fig. S4**). This work further demonstrates the remarkable conservation of transmembrane domains forming the MDMA and serotonin binding pocket in octopus SLC6A4 paralogues, including complete conservation of amino acids implicated in MDMA binding in human.

**Fig. 3.**
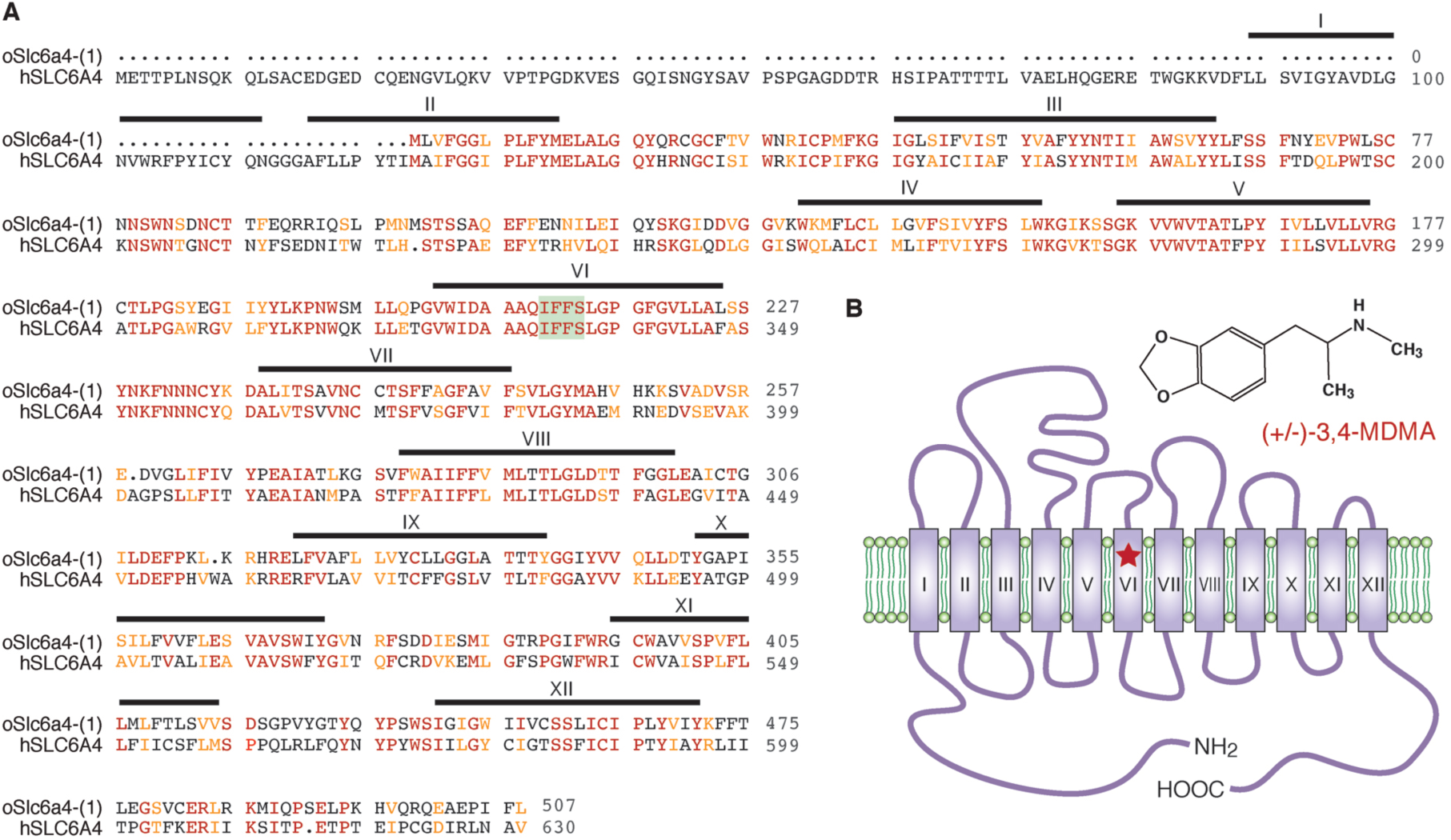
Protein alignment of *O. bimaculoides* Slc6a4-(1) to human SLC6A4. **A.** Pairwise protein alignment. Human SLC6A4 transmembrane domain 1-12 annotations are based on Uniprot Topology annotation of protein P31645 (SC6A4_HUMAN). The binding pocket region that overlaps serotonin and MDMA binding (green) based on (*19*). Alignment of annotated human SLC6A4 to octopus Slc6a4-(1) was done by pairwise alignment in Geneious. **B.** Diagram illustrating the structure of MDMA, transmembrane domains of the SLC6A4 protein, and the MDMA binding site (red star).

Encouraged by these results, we next sought to test the functional conservation of MDMA’s effects in *O. bimaculoides.* As diagramed in **Fig. 4A-B**, baseline social approach behaviors were tested in drug naïve subjects in an arena containing a novel object and a male social object for 30 minutes (pre trial). Two to 24 hours later, subject animals were placed in a bath containing MDMA for 10 minutes, followed by a 20-minute saline wash, and then tested again for 30 minutes (post trial). As shown in **Fig. 4C**, during the pre trial subject animals spent significantly more time in the object chamber compared to the center chamber, whereas in the post trial, subject animals spent significantly more time in the social chamber compared to the center chamber. Comparisons between pre versus post MDMA conditions revealed that the time spent with the social object was significantly increased following MDMA treatment (**Fig. 4H-I**), while the time spent in the object and center chambers was not significantly different across conditions (**Fig. 4D-G**). Quantification of the number of transitions between chambers indicated no significant increase in locomotor activity (**Fig. 4J-K**), nevertheless, qualitatively, in all subjects we did observe slow, voluntary, non-stereotypical, semi-purposeful movements following MDMA treatment. Together these results demonstrate that the acute prosocial effects of MDMA are conserved in *O. bimaculoides*, and suggest that this pharmacological manipulation releases extant, but normally suppressed, neural mechanisms subserving social behaviors.

**Fig. 4.**
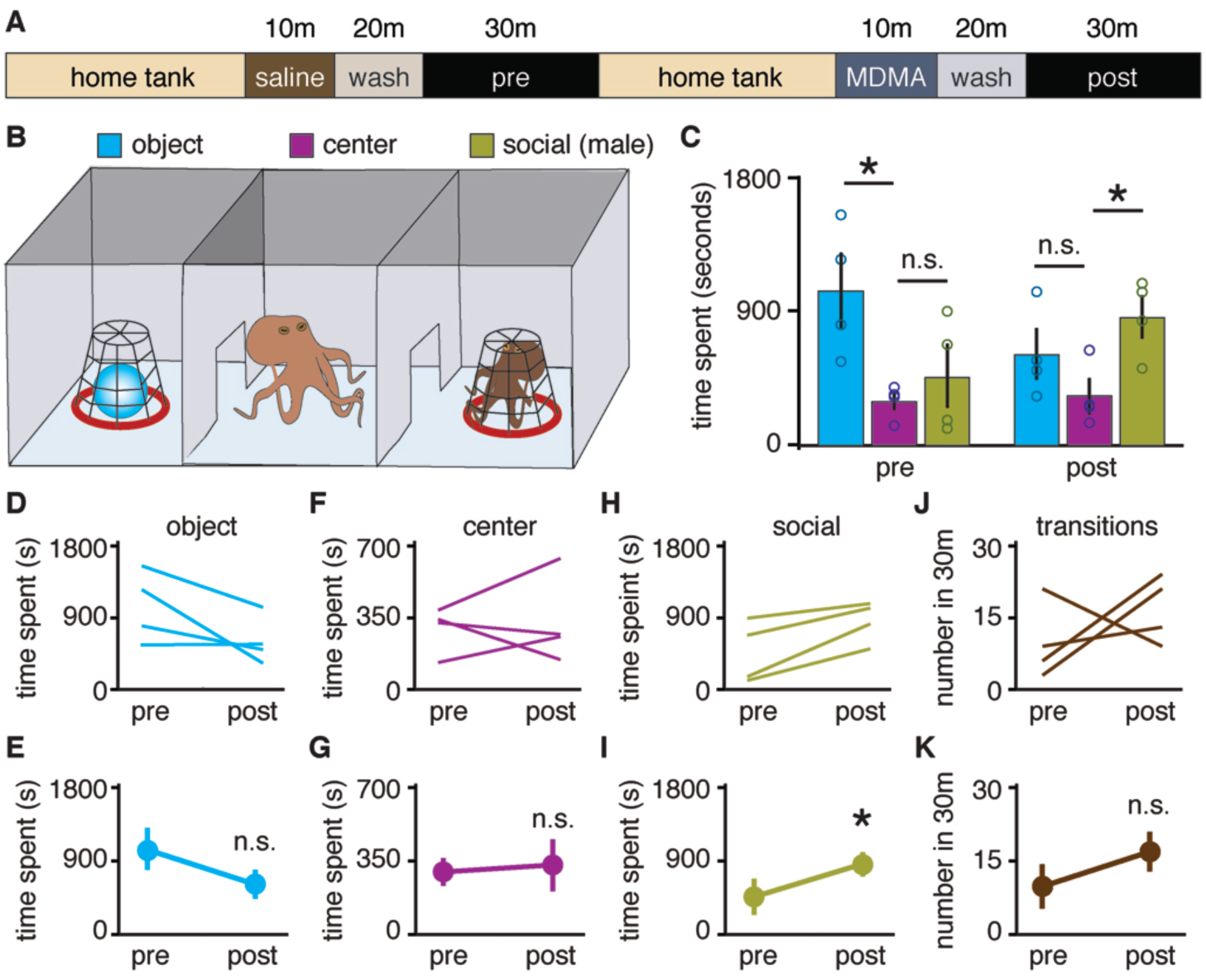
Prosocial effects of MDMA in *O. bimaculoides*. **A-B** Diagrams illustrating timeline (A) and experimental protocol (B) for three-chambered social approach assay. **C**. Quantification of time spent in each chambers during 30-minute test sessions (n = 4, Two-way repeated measures ANOVA: P = 0.0157; post-hoc unpaired t-test pre, social versus center P = 0.4301, object versus center P = 0.0175; post, social versus center P = 0.0190, object versus center P = 0.1781). **D-K**. Comparisons between pre versus post MDMA treatment conditions. Paired t-test pre versus post, social time P = 0.0274; object time P = 0.1139; center time P = 0.7658; transitions P = 0.3993).

The current studies are the first to experimentally quantify social approach behaviors in *O. bimaculoides* (**Fig. 1A-K**), and demonstrate that consistent with previous ethological descriptions (*6*), this species shows no preference for social approach to a novel male conspecific (**Fig. 1A-H**). Nevertheless, somewhat surprisingly, both male and female subjects did exhibit social approach to a novel female conspecific (**Fig. 1A-H**), a finding that may reflect an adaptation of laboratory raised animals, or an incomplete ethological description of the full repertoire of social behaviors in the wild (*12*). At the same time, unlike mice (*10*), social novelty was not a superseding factor in determining social approach behaviors in *O. bimaculoides* (**Fig. 1I-K**). Bioinformatics studies revealed clear orthologs of human SLC6A4 in octopus, as well as high levels of conservation in the transmembrane domain and amino acid region critical to MDMA binding (*19*). Interestingly, we found that SLC6A4 is broadly conserved in fruit fly, worm, and most other bilaterian animals, but is surprisingly absent in both of the eusocial hymenopteran insects, honey bee and leaf cutter ant. This absence raises the possibility that in these eusocial invertebrates, sociality evolved convergently utilizing other neurotransmitters or peptide hormones (*24*, *25*). The current studies also provide the first functional evidence that the prosocial effects of MDMA (*26*) are evolutionarily conserved in *O. bimaculoides* (**Fig. 4**). Although we did not observe a significant increase in locomotor activity following MDMA, this finding is consistent with the complex locomotor profile of MDMA reported in rodents (*17*, *27*).

Based on our ability to induce prosocial approach behaviors by manipulating serotonergic signaling, it is tempting to speculate that in octopuses sociality is the default state, which is suppressed outside ethologically relevant periods, such as mating. Additionally, the current studies establish the first drug delivery protocols for pharmacological experiments in octopus, and indicate that effective doses of MDMA are in the same range as those described for humans and rodents. Beyond their utility for the current studies, development of this experimental infrastructure fulfills an unmet need for the field, and will enable future mechanistic studies in octopuses. Moreover, this work demonstrates that medications currently in use or under investigation in humans (*22*, *28*–*30*), target a homologous binding site in *O. bimaculoides*, and provides important proof of concept data supporting further development of octopuses as model organisms for translational research.

**Full Methods** and any associated references are available in the online version of the paper.

## Acknowledgements.

We thank members of the Dölen laboratory, R. Caldwell, M. Kuba, and T. Gutnick for comments, as well as K. Peramba, M. Renard, K. Dever, D. Calzarette, L. Menlow, J. Simmons, J. Marvel-Zuccola, J. Miao, M. Cordeiro, P. Newstein, and T. Sakmar for guidance and assistance in octopus care, and D. Mark Welch and M. Sogin for guidance in the phylogenetic analysis. MDMA was a gift of R. Doblin (Multidisciplinary Association For Psychedelic studies, MAPS). This work was supported by grants from the Kinship Foundation, Hartwell Foundation, and Klingenstein-Simons Foundation (G.D.) and the Vetlesen Foundation (E.E.)

## Author Contributions

G.D. and E.E. designed the study, interpreted results and wrote the paper. G.D. and E.E. performed behavioral and pharmacological experiments. E.E. performed bioinformatics analysis and octopus culturing. G.D. performed statistical analysis. All authors edited the paper.

## Supplementary Materials

**Fig. S1. Consensus bootstrap tree for the SLC6A4 gene family.**

**Fig. S2. SLC6A monoamine transporters subtree to highlight species composition per clade and branch lengths.**

**Fig. S3. Protein alignment of *O. bimaculoides* Slc6a4-(2) to human SLC6A4.**

**Fig. S4. Details of protein and domain alignments for *O. bimaculoides* Slc6a4-(1) and Slc6a4-(2) to human SLC6A4.**

**Fig. S5. Protein alignment of mollusc SLC6A4 proteins to human SLC6A4.**

**Fig. S6. A larger version of Fig. 1A.**

**Table S1. SLC6A reference gene set for phylogenetic analysis.**

**Table S2. Species and proteome sources for phylogenetic analysis.**

**Table S3. Mapping of species to their SLC6A4 gene and tree identifiers**

